# Electrophysiological signatures of brain aging in autism spectrum disorder

**DOI:** 10.1101/2021.03.18.434885

**Authors:** Abigail Dickinson, Shafali Jeste, Elizabeth Milne

## Abstract

Emerging evidence suggests that aging processes may be altered in adults with autism spectrum disorder (ASD). However, it remains unclear if oscillatory slowing, a key neurophysiological change in the aging brain, manifests atypically in this population. This study sought to examine patterns of age-related oscillatory slowing in adults with ASD, captured by reductions in the brain’s peak alpha frequency. Resting-state EEG data from adults (18-70 years) with ASD (N=93) and age-matched neurotypical (NT) controls (N=87) were pooled from three independent datasets. A robust curve-fitting procedure quantified the peak frequency of alpha oscillations (7-13Hz) across all brain regions. Associations between peak alpha frequency and age were assessed and compared between groups. Consistent with characteristic patterns of oscillatory slowing, peak alpha frequency was negatively associated with age across the entire sample (p<.0001). A significant group by age interaction revealed that this relationship was more pronounced in adults with ASD (p<.01), suggesting that that age-related oscillatory slowing may be accelerated in this population. Scalable EEG measures such as peak alpha frequency could provide insights into neural aging that are crucially needed to inform care plans and preventive interventions that can promote successful aging in ASD.

## 1. Introduction

Biological aging processes alter neuronal and synaptic function in later life.^1,2^ As a result, oscillatory brain dynamics become slower, captured by reductions in the brain’s peak alpha frequency (7-13Hz).^3–6^ These changes undermine the temporal framework through which neuronal populations communicate and are associated with age-related deterioration of long-range functional connectivity ^7–11^ and cognitive functions that depend upon distributed processing,^12–14^ such as executive function, memory, and processing speed.^15^ As such, patterns of brain activity provide insight into trajectories of cognitive decline, which can vastly differ between individuals of the same age. For instance, steeper reductions in peak alpha frequency and functional connectivity predict more significant deficits in memory, attention, and decision-making, ^14–17^ suggesting that the pace of oscillatory aging varies across the general population and may contribute to individual differences in cognitive decline. However, while these studies have shed light on typical levels of variability, it is unclear how aging processes impact adults with pre-existing neural differences. Age-related changes may interact with neuropathology, rendering patterns of brain activity more vulnerable to disruption.

The present study examines neural aging in autism spectrum disorder (ASD), a lifelong neurodevelopmental condition characterized by social communication impairments and restricted, repetitive behaviors ^18^. Despite affecting more than 2% of American adults, a lack of lifespan research has severely limited our understanding of aging processes in ASD. As a result, specific mechanisms and risk factors associated with aging remain undetermined in this population. Recent cognitive and neuroimaging evidence consistent with atypical brain aging in ASD emphasizes the critical need to bridge this knowledge gap. Compared to the general population, adults with ASD exhibit more significant declines in executive function skills ^19,20^ and experience higher rates of cognitive disorders. ^21^ Consistent with these phenotypic changes, magnetic resonance imaging (MRI) studies report steeper white matter deterioration ^22^ and accelerated functional connectivity reductions <u>(1,2)</u> in adults aging with ASD. However, it is unclear if neurophysiological aging mechanisms are altered in this population.

As a scalable and sensitive assay of neurophysiological aging, electroencephalography (EEG) measures of peak alpha frequency hold promise for detecting early neural deviations that precede behavioral declines. In turn, mapping patterns of oscillatory aging in ASD could reveal pathways of risk and resilience that inform opportunities for targeted interventions to improve aging outcomes in this vulnerable population. In line with these overarching goals, the present study characterizes the relationship between age and EEG measures of peak alpha frequency in ASD, examining whether patterns of cross-sectional change differ from those of age-matched controls and if characteristics such as symptom severity or cognitive function influence are associated with peak alpha frequency.

## 2. Materials and Methods

### 2.1. Participants

This study examined EEG recordings from 180 adult participants. Ninety-three participants (26 female) had previously received an ASD diagnosis from a trained clinician according to DSM-IV, DSM-V or ICD-10 criteria. Eighty-seven participants (34 female) without ASD, nor any first-degree relatives with ASD, comprised a ‘neurotypical’ (NT) control group. Data for this sample were obtained from three independent sources. Dataset 1 was acquired from a previously published EEG study ^24^, wherein participants consented to data sharing for secondary analysis. Datasets 2 and 3 were obtained from the National Institute of Mental Health (NIMH) Data Archive (NDA). Detailed protocols for each dataset are publicly available (see acknowledgment section) and described in Table 1. Full details regarding the criteria used to acquire data from NDA are described in the supplementary materials.

**Table 1.**
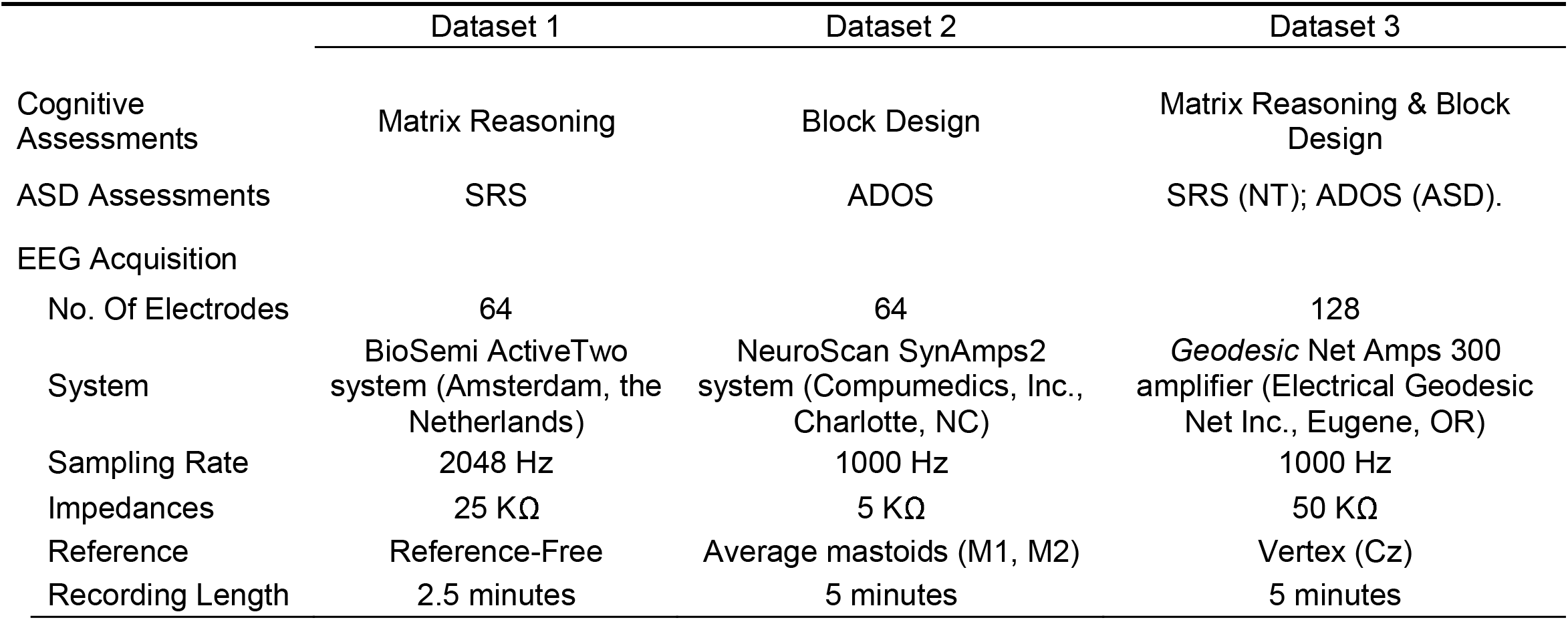
Study protocol details for each dataset.

The age and sex of ASD and NT groups did not significantly differ. Consistent with their clinical diagnosis, adults with ASD had increased social communication difficulties and decreased cognitive abilities compared to NT adults. Demographic information are described in Tables 2. Supplementary table S2 details demographic information for each separate cohort, and table S2 describes concomitant conditions and psychoactive medications at the time of EEG recording for both ASD and NT groups.

**Table 2.** Participant characteristics.

| | ASD<br>(n=93) | NT<br>(n=87) | Group Comparison<br>Student's *t* or $X^2$<br>*p* value |
| --- | --- | --- | --- |
| Age (Years) |  |  |  |
| Mean (SD) | 30.39 (13.64) | 32.52 (12.30) | .274 |
| Range | 18.00-67 | 18-69.00 |  |
| Sex |  |  |  |
| N females (% Female) | 26 (28%) | 34 (39.1%) | .118 |
| NVIQ |  |  |  |
| *n* completed | 89 | 85 |  |
| Mean (SD) | 52.64 (10.21) | 57.16 (9.20) | .003 |
| Range | 26-72 | 33-77 |  |
| SRS |  |  |  |
| *n* completed | 28 | 52 |  |
| Mean (SD) | 67.71 (10.01) | 45.32 (5.62) | <.001 |
| Range | 48-84 | 38-62 |  |
| ADOS |  |  |  |
| *n* completed | 64 | 34 |  |
| Mean (SD) | 10.61 (3.49) | 2.18 (2.21) | <.001 |
| Range | 4-21 | 0-7 |  |

Note. ASD symptoms were assessed using either SRS or ADOS. There was no overlap between participants who completed (described as 'n completed' in Table 1) either of these assessments.

### 2.2. Behavioral Assessments

#### 2.2.1. Cognitive Function

Non-verbal cognitive function was quantified using scores from the Wechsler Abbreviated Scale of Intelligence (WASI) ^25^, a standardized measure of cognitive skills that includes four subtests (vocabulary, similarities, block design [BD], and matrix reasoning [MR]). Non-verbal IQ scores (NVIQ) were calculated for Dataset 3 by averaging the subscale T-scores, which take age into account, from the BD and MR subscales. As only one subscale was available for Datasets 1 (MR) and 2 (BD), these scores provided a proxy for NVIQ. Measures of verbal cognitive function were not collected in Dataset 1. As such, this study did not examine verbal IQ. However, non-verbal cognition may represent a less biased estimate of underlying cognitive differences between the ASD and NT groups, given that verbal IQ measures can be influenced by the core communication and language deficits that characterize ASD.^26^ Cognitive assessments were not available for 6 participants (ASD: 4, NT: 2). Additional behavioral assessments confirmed diagnoses in the six participants with ASD who scored <60 on the SRS (See Supplementary Materials).

#### 2.2.2. ASD Symptoms

ASD symptoms and social communication abilities were quantified using calibrated severity scores (CSS) from module four of the Autism Diagnostic Observation Schedule (ADOS, ^27^) and the adult self-report version of the Social Responsiveness Scale 2nd Edition (SRS-2;^28^), respectively. The ADOS is an assessment tool used by clinicians and researchers to assess social-communication and repetitive behaviors associated with ASD. Higher CSS scores indicate increased symptom severity, with a CSS ≥ 7 consistent with a clinically relevant level of ASD symptoms. The SRS-2 is a 65-item questionnaire, with higher scores indicating reciprocal social communication difficulties. Normed SRS T-scores over 60 indicate reciprocal social behavior deficiencies. Six participants with ASD scored <60 on the SRS. As a result, additional behavioral assessments confirmed ASD diagnoses in these participants (see supplementary material for detailed description). As expected, SRS and ADOS scores were significantly higher in participants with ASD (p<.001). Information regarding ASD symptoms were not available for two participants (ASD: 1, NT: 1).

### 2.3. EEG acquisition & Processing

Each participant contributed >2.5 minutes of eyes-closed resting EEG data acquired using high-density research systems (see Table 1). In-house MATLAB scripts and EEGLAB ^29^ were used to implement a data processing pipeline (described below) that was specifically designed to harmonize EEG data across different systems, facilitating the pooling of three independent datasets. Full details regarding data harmonization can be found in the supplementary material.

Data were high-pass filtered to remove frequencies below 1 Hz and resampled to 512Hz. Continuous data were then visually inspected, and sections of data showing excessive electromyogram (EMG) or other non-stereotyped artifacts were removed. Data were interpolated to the international 10-20 system 25 channel montage ^30^ to facilitate comparisons with previous studies of PAF in children with ASD ^31^. Independent component analysis (ICA), a statistical blind source separation technique, was used to decompose data into maximally independent components (IC) ^32^. Any IC that represented stereotyped artifacts (including EMG, EOG, heart artifact, and line noise) were removed from the data using the eeglab function iclabel (components <75% neural). Artifact subspace reconstruction (ASR), a data cleaning method that uses sliding window principal component analysis, was then used to remove high amplitude artifacts, relative to artifact-free reference data ^33,34^. The eeglab function clean_RawData implemented ASR, with default parameters and rejection threshold k=20 ^33^. The first 120 seconds of data for each participant (which represented the minimum amount of cleaned data available across participants) underwent further analyses.

#### 2.3.1. Peak alpha frequency (PAF)

Continuous EEG data were transformed into the frequency domain using Welch’s method (1024 sample Hamming-tapered windows with 50% overlap), resulting in power spectral density estimates with 0.5 Hz frequency resolution. Average power spectral density estimates (averaged over all channels) are displayed in (Fig 2A). Analyses focused on the computed power spectra of six channels (F3, F4, C3, C4, O1, and O2) across three regions of interest (frontal, central & occipital), with each region defined by the average of two channels (F3, F4 = frontal; C3, C4 = central; O1, O2 = occipital). The 1/f trend of the log-transformed power spectrum (1–55 Hz) of each channel was first modelled using a least-squares method and then subtracted from the data ^35,36^, facilitating peak identification without bias towards lower frequencies ^37^. De-trended spectra are displayed in (Fig 2B). Using a robust curve-fitting procedure, we then selected the peak of a Gaussian curve fit within the alpha range (7-13Hz) to quantify PAF, thus avoiding the ambiguity of choosing the local maxima. See Dickinson et al., 2017 for a detailed description. This slightly larger alpha range of 7-13Hz accommodated peak identification across all participants, including those who may have decreased PAF due to advanced age. Representative peak examples are displayed in (Fig 2C). If the curve-fitting procedure failed for de-trended spectra, this indicated a lack of modulation within the alpha range, and no peak was selected. Four participants did not show an alpha peak in a single brain region (3 ASD; 1 NT): one within the frontal region, two within the central region, and one within the occipital region. The number of missing PAF values did not differ between ASD and NT groups.

**Figure 1.**
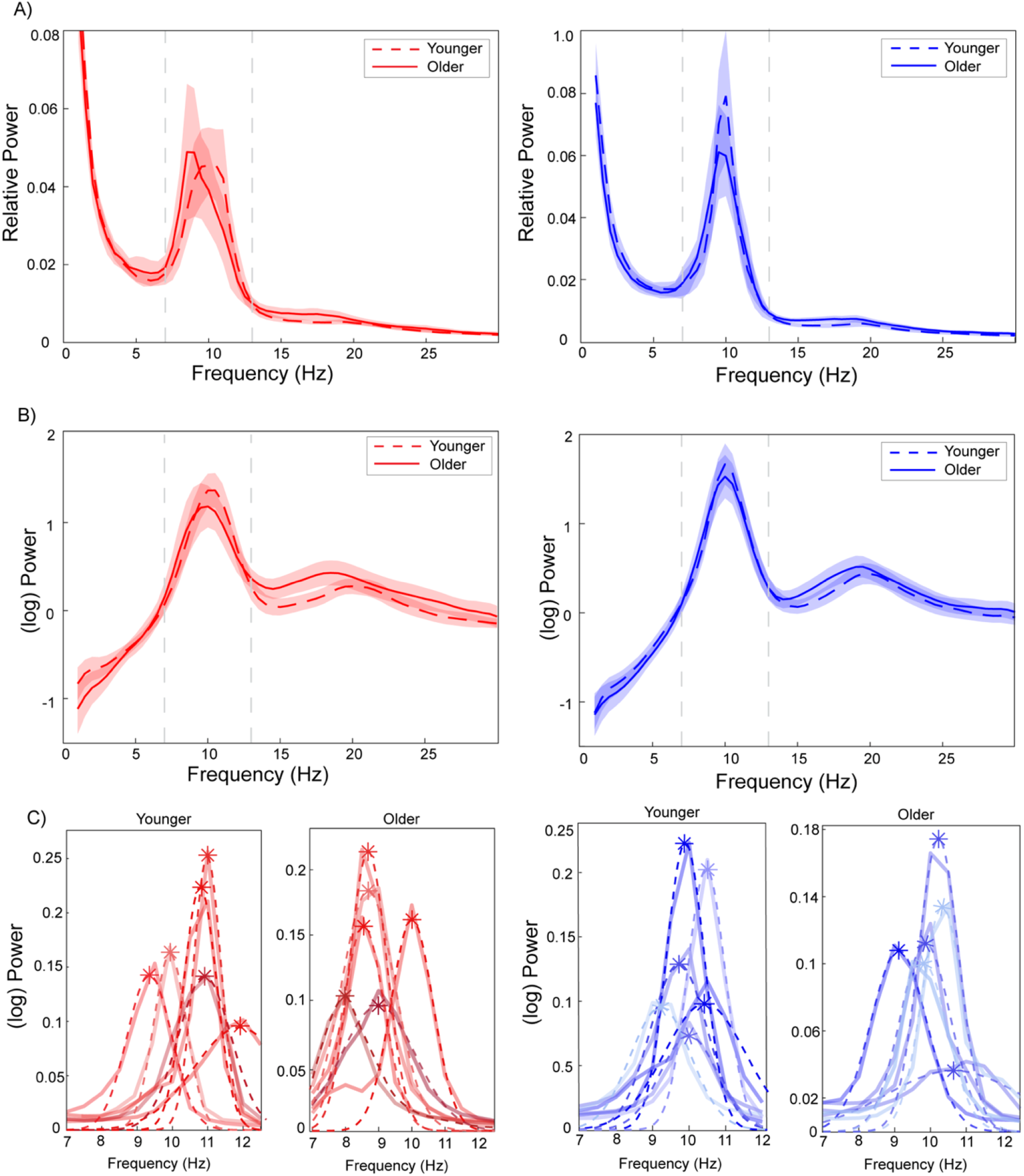
Example of the PAF fitting routine across all ASD participants (red, left panel), and NT participants (blue, right panel). (A) Log-transformed power spectral density (average of three regions of interest) for younger (dashed line) and older adults (solid line). Shaded areas represent 95% confidence intervals. (B) Log-detrended power spectra for younger (dashed line) and older adults (solid line). Shaded areas represent 95% confidence intervals. (C). Examples of representative individual power spectra with fitted Gaussian curves for several younger and older participants in each diagnostic group. Different shades of red and blue represent individual participants. The dashed line represents the Gaussian curve fit, with asterisks denoting the peak alpha frequency value.

**Figure 2.**
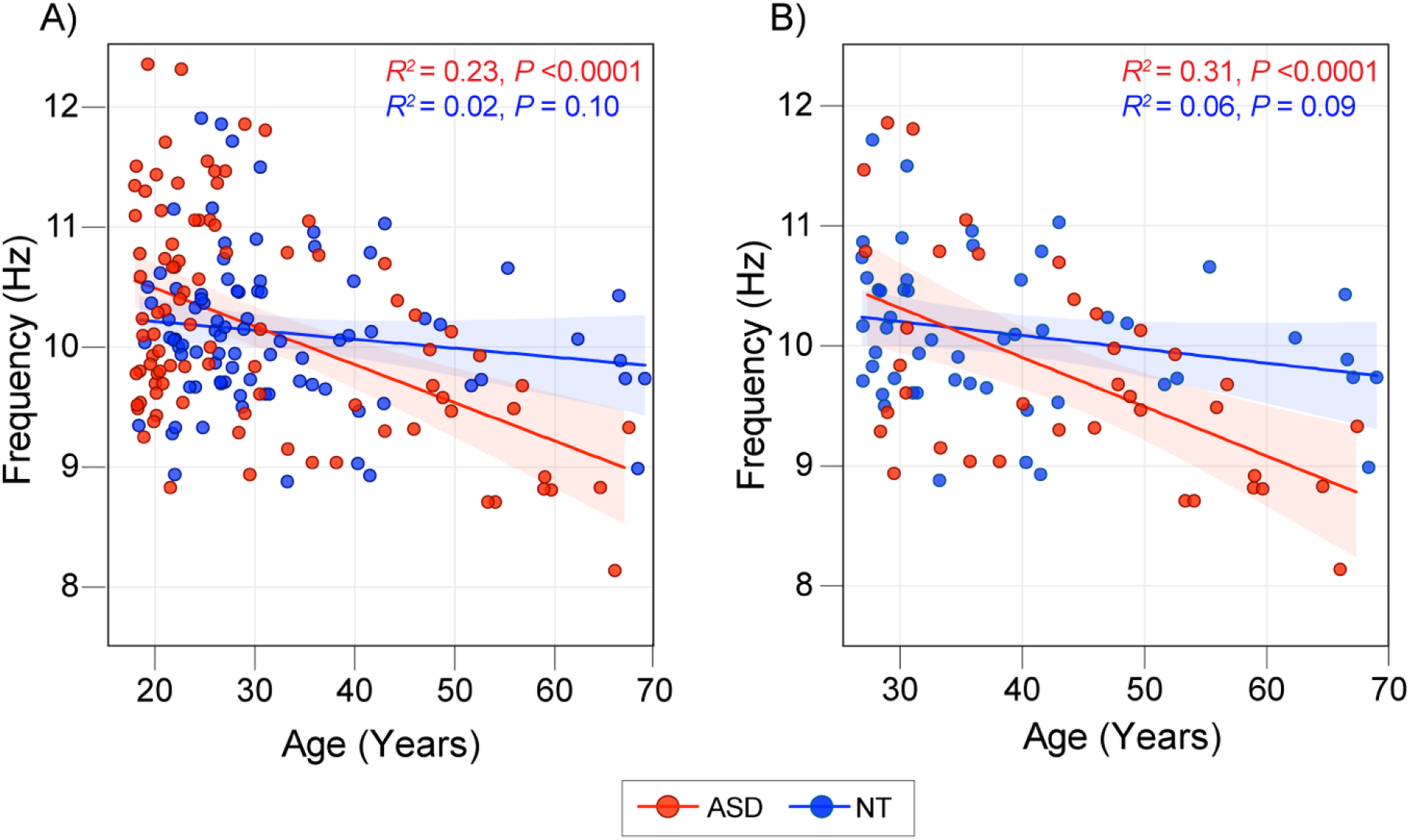
Differential association between age and peak alpha frequency according to ASD diagnosis in (A)The full sample and (B) in participants over 27 years of age. Shaded areas represent 95% confidence intervals. Peak alpha frequency values are marginal means after controlling for covariates in the baseline model.

### 2.4. Statistics

Hierarchical linear regression analyses examined associations between PAF and age. To ensure that study protocol (i.e., dataset number), gender, or NVIQ did not confound estimates of PAF decline, these variables were entered in the first block of the hierarchical regression model. Age, group (ASD or NT), and group by age interaction terms were then entered as predictors of PAF. Significant interactions reflecting group differences were further explored by comparing Fisher r-to-z transformed correlation coefficients between the two groups using an independent samples t-test. Given the known trajectory of PAF decline ^38^, linear relationships were examined a priori. We conducted additional analyses to determine if the relationship between age and PAF within each group was better fit by a nonlinear model with age entered as a quadratic term. Model performance was assessed using the corrected Akaike information criterion (AICc).

Linear regressions controlling for the increased presence of medication and concomitant clinical diagnoses in the ASD group followed up relationships between age and PAF. To account for the over-representation of younger participants in both the ASD and NT groups, we also repeated analyses solely considering older participants (>27 years, the sample’s median age).

Hierarchical linear regressions examined the relationship between cognitive function and PAF, with non-verbal cognition (NVIQ) entered as the dependent variable and group (ASD or NT), PAF, and the interaction between PAF and group entered as predictors. Similar to the hierarchical regression described above, a baseline model accounted for the confounding effects of age, sex, and study protocol. Finally, exploratory regressions analyzed the relationship between ASD symptom severity (ADOS and SRS scores) and PAF in ASD and NT groups separately, with sex, age, and study protocol entered as covariates.

## 3. Results

### 3.1. Preliminary Analyses

Preliminary analyses considered regional differences in PAF decline using a repeated-measures ANCOVA. There was no significant interaction between region and age (*F*_2, 344_ =.039, *p*=.672), nor between group, region, and age (*F*_2, 344_=.042, *p*=.733), indicating that age was similarly associated with PAF across all regions of interest. We therefore collapsed PAF across regions for all further analyses (see Table 3 for regional PAF values). Average PAF did not show sex differences (*t*_178_=-.129, *p*=.89), nor differences across the three independent datasets (*F*_2, 177_=.867, *p*=.42).

**Table 3.**
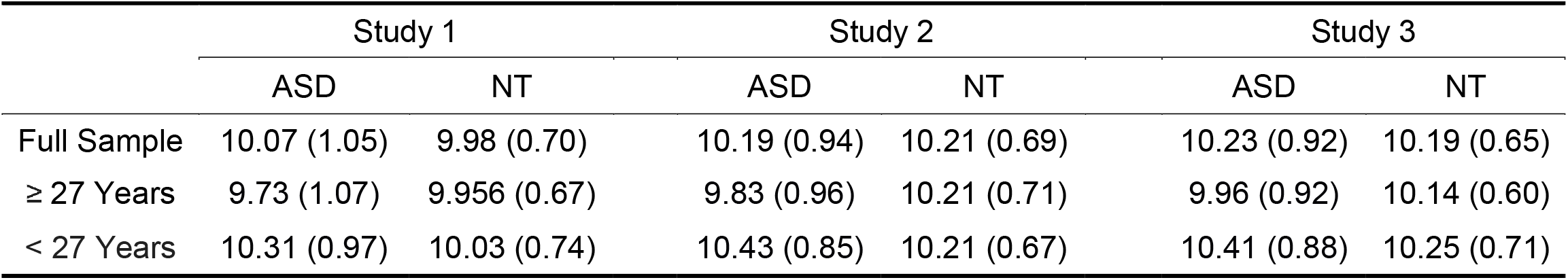
Peak alpha frequency values (Mean(SD)) for each region of interest.

### 3.2. PAF and Age

Age was significantly associated with PAF across all participants (*F*_4, 171_=6.10, *p*<.0001). A significant group by age interaction (*F*_6, 165_=5.435, *p*=.01) was due to a stronger (*z*= -2.168, *p*= 0.015) relationship between age and PAF in the ASD group (*r* =-.443, *p* < .001) than in the NT group (*r* =-.1468, *p*=.09; Fig 3A). AIC values indicated that the relationship between PAF and age was better fit by a nonlinear model in the ASD group (Δ AICc= - 1.196) and a linear model in the NT group (Δ AICc= 1.795). The full regression table is provided in the supplementary material (Table S3).

**Figure 3.**
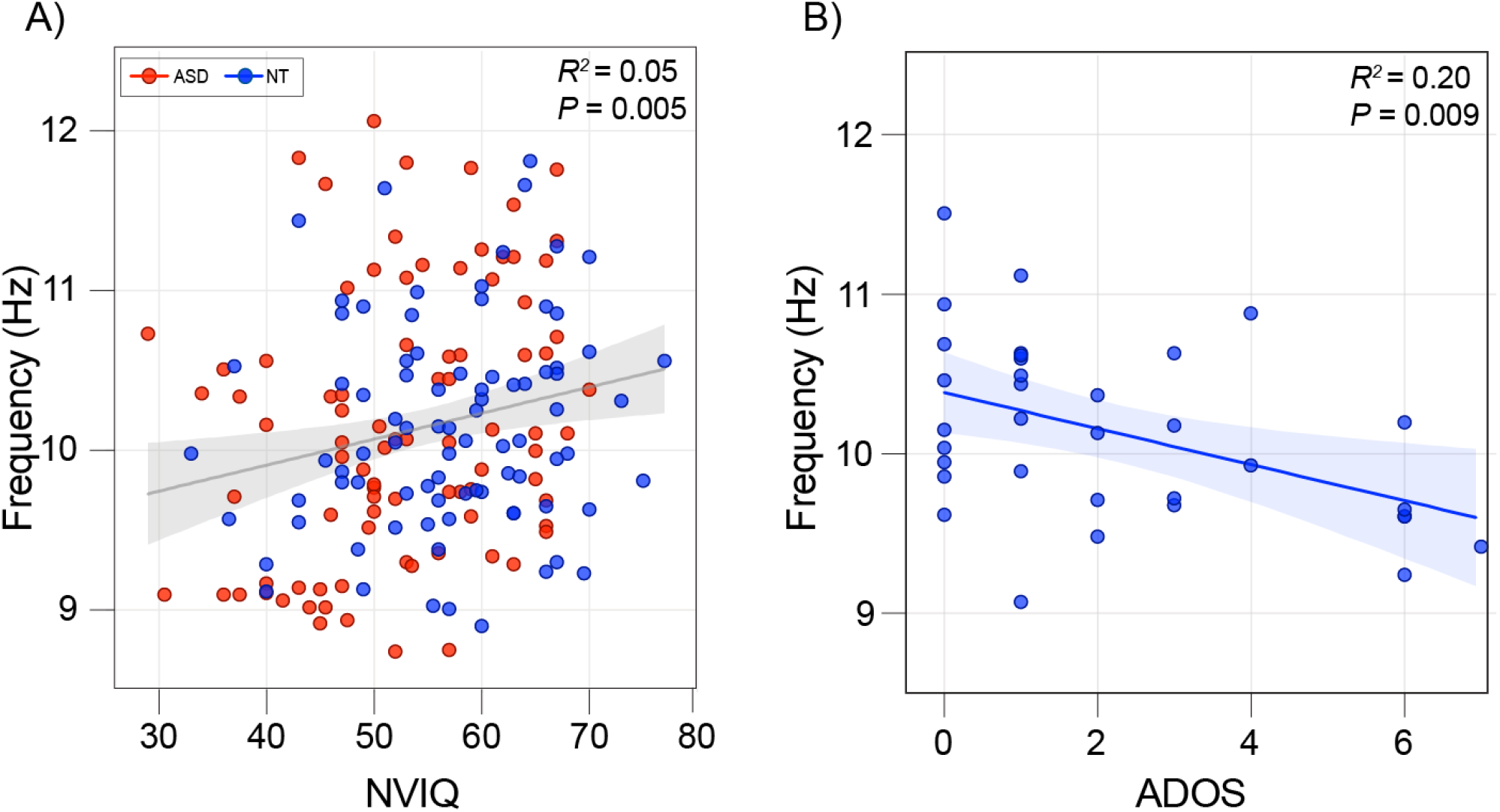
Associations between peak alpha frequency are behavioral measures. (A) Peak alpha frequency was associated with NVIQ across the entire sample. (B) Within NT adults, peak alpha frequency was associated with ADOS score. Shaded areas represent 95% confidence intervals.

Additional analyses confirmed that PAF did not differ according to the presence of medication (*p*=.83) or concomitant diagnoses (*p*=.68), indicating that these factors did not drive group differences in the relationship between age and PAF. Further, the association between age and PAF (*p*<0.00001), as well as the interaction between group and age (*p*=.007), also remained significant after controlling for medication and concomitant diagnoses by adding them into the baseline model. Finally, the group by age interaction remained significant when only considering older participants (*F*_1,86_=4.59, *p*=.023) due to a stronger association between age and PAF in the ASD group (*r*=-.582, *p*<.001) compared to the NT group (*r*=-.248, *p*=.041; Fig 3B).

### 3.3. PAF and Behavioral Measures

While NVIQ was a significant predictor of PAF (*F*_1, 169_=7.56, *p*<.007), there was no significant group interaction (P=.894), indicating that the positive correlation between PAF and NVIQ (*r*=.177, *p*=.02) was consistent across participants, regardless of diagnostic status. The relationship between NVIQ and PAF also remained significant when controlling for age (*r*=.207, *p*=<.007; Fig 4A).

**Figure 4.**
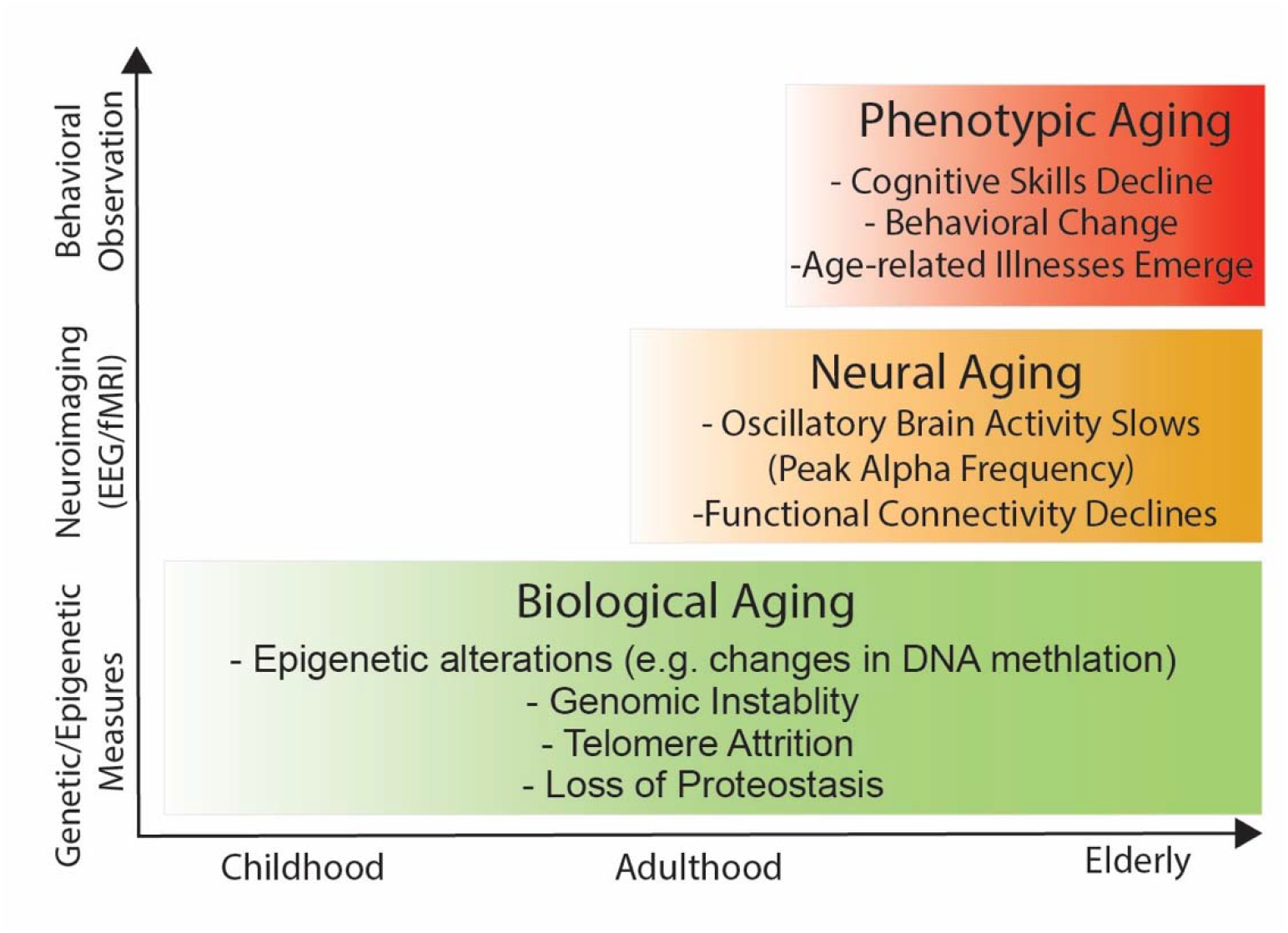
Aging is a complex, non-linear process that involves progressive changes across several scales of measurement in the brain. On a molecular level, ‘hallmark’ biological changes cause neurobiological changes and cellular damage that eventually disrupts large-scale brain function. In turn, changes in network-level communication disrupt cognitive processes such as memory and processing speed. Note that peak alpha frequency (PAF decline) represents a network-level change.

Exploratory analyses considered the association between behavioral ASD assessments and PAF in each group separately. In the ASD group, ASD symptoms (quantified by ADOS scores) and social communication difficulties (quantified by SRS scores) were not associated with PAF (*p*>.05). In the NT group, social communication difficulties were not associated with PAF (*p*>.05; *n*=52). However, ASD symptoms were significantly associated with PAF (*r*=-.447, *p*=.008; *n*=34). This relationship remained significant after controlling for age and NVIQ (*r*=-.431, *p*=.017; Fig 4B). As described in the methods section, there was no overlap between NT participants who received the SRS and those who received the ADOS (see table), which may account for the divergent relationships seen across the two measures.

## 4. Discussion

This study demonstrates cross-sectional differences in the association between age and PAF in adults with ASD. Specifically, the quadratic PAF decline of ∼ 3.3% per decade in ASD was significantly amplified compared to the linear decline of ∼0.7% per decade in NT adults. A steeper rate of change suggests that age-related oscillatory slowing may be accelerated, or even premature, in adults with ASD. In contrast, the linear age-related decline observed in NT participants follows a similar trajectory to previous reports in larger samples ^38^, suggesting that attenuated PAF changes in the NT group did not drive the present findings. Finally, the present results replicate a well-established positive association between PAF and non-verbal cognitive abilities across both groups ^4,6,39–41^, indicating that PAF captures oscillatory characteristics associated with cognitive processing.

Overall, the present results are consistent with an atypical trajectory of neurophysiological aging in ASD, aligning with other recent evidence of earlier, or more rapid, age-related neural changes in adults with ASD. Our findings are particularly relevant to reports of accelerated white matter deterioration in ASD, given that PAF shows a close association with white matter architecture in NT populations ^42,43^ and captures typical age-related declines in white matter integrity ^44–46^. Along with mounting evidence that adults with ASD are at greater risk of cognitive decline ^20^ and age-related cognitive disorders, ^21^ these data reinforce a crucial need for investigations into mechanisms of aging in ASD. Studying aging pathways and related risk factors that contribute to these disparities is particularly urgent, given that more than a million American adults with ASD will be over 65 years of age by 2035, outnumbering children with ASD for the first time.^47,48^

As a marker of neurophysiology, PAF holds promise for capturing the cascading and complex processes that affect the aging brain across multiple levels of neural organization. Figure 4 illustrates how intermediate neurophysiological processes reflect variation in underlying biological aging mechanisms and shed light on downstream phenotypic alterations, including cognitive decline. As such, PAF could provide a crucial opportunity to detect early neurophysiological deviations before cognitive declines begin to impact quality of life in ASD ^49,50^. It should be noted that this opportunity is not exclusive to ASD, reinforced by our finding that the association between PAF and cognitive measures is present within both groups, as well as previous studies linking PAF and cognitive function in other populations. ^3114,51^ However, earlier predictive markers of cognitive decline are especially needed in ASD, given recent evidence of disproportionate age-related declines in executive function and daily living skills. ^20,52^

PAF may also provide insight into the underlying mechanisms that drive atypical changes in brain function and structure in ASD, helping to disentangle 1) why these processes may be altered, and 2) how they map onto cognitive and behavioral changes. One possibility is that neural aging processes interact with pre-existing neuropathology in ASD, making adults with ASD more sensitive to typical levels of age-related neural deterioration. Fundamental processes during early brain development are thought to be altered in ASD, disrupting typical circuit formation and altering oscillatory communication between higher-order brain regions.^53–56^ A vast literature describes atypical neurophysiology in ASD, including the atypical development of peak alpha frequency during childhood ^57,58^ and atypical long-range oscillatory coherence. ^59–62^ Given these differences stemming from early brain development, adults with ASD may be more vulnerable to further neural changes that occur due to typical biological aging processes. Evidence from neurotypical populations supports the possibility that that pre-existing neural differences could increase vulnerability for later age-related changes. For instance, one functional magnetic resonance imaging (fMRI) study reports that baseline reductions in large-scale connectivity predict more rapid functional declines in healthy adults ^63^. These data suggest that pre-existing differences may render brain networks in ASD more sensitive to typical levels of age-related deterioration. Studying how patterns of brain activity change longitudinally could help shed light on this possibility and potentially elucidate the cellular and molecular variations that alter their dynamics.

Studying neural aging could also help to identify if the ASD phenotype itself may alter outcomes, leads. It is well-established that the social environment can influence epigenetic aging mechanisms, with environmental factors accounting for much larger variations in human longevity (>70%) than genetic factors, which contribute 20-30%.^64–66^ In the context of typical aging, social isolation and loneliness increase risk of mortality ^67^ and age-related health problems, such as cardiovascular disease, chronic respiratory diseases, cancer, diabetes, and dementia. ^68–70^ Social isolation is also linked to accelerated brain aging, demonstrated by declines in global cognition^68,71–74^ and reductions in functional connectivity^75^ and peak alpha frequency.^76^ However, the link between social isolation and aging outcomes has not been investigated in ASD. This question is particularly relevant when considering that adults with ASD interact less with friends, participate in fewer social activities, and experience increased loneliness compared to non-ASD peers.^77,78^

Neurophysiological markers could provide objective insight into how social differences impact neural aging processes and how social impairments in ASD contribute to this relationship. With these goals in mind, the present study conducted exploratory analyses of the associations between social behaviors and PAF. While the present results do not find a significant association between symptom severity and PAF within the ASD group, sub-clinical ASD symptoms in the NT group (measured using ADOS) were negatively associated with PAF, possibly reflecting an association between social connectedness and neurophysiological aging. However, this finding has several limitations, as it only represents the small subset of NT participants assessed using the ADOS. The subset who received the SRS, a more appropriate measure of social communication in NT adults, did not show a similar association between social measures and PAF. Future research with comprehensive assessments of social impairments and social integration measures is needed to investigate this potential relationship in a broader ASD population.

In addition to examining the mechanistic pathways that influence aging outcomes in ASD, a better understanding of neural aging in ASD may highlight factors that place individuals at higher risk of sub-optimal trajectories and inform pathways of risk and resilience to improve aging outcomes in ASD. As a scalable technique that could monitor functional brain changes across large, diverse populations, EEG is particularly well-suited for this endeavor and would have broad applicability for the early identification of accelerated neurophysiological aging in ASD and other neurodevelopmental disease processes. In turn, mapping age-related changes at an individual level may highlight risk factors that could be modified through preemptive interventions that begin early in life. These efforts could include reducing social isolation in ASD through other sources of social contact shown to improve aging outcomes in the general population, including pets,^79^ online interactions,^80^ and videogames. ^81^ Other protective factors that could be promoted in at-risk populations include increasing resilience ^82^, and managing co-morbid medical conditions that hold independent age-related risks ^83^.

Finally, as a marker of potential neurophysiological aging differences in ASD, PAF could also inform targets for future biological treatment modalities and help evaluate interventions that target these mechanisms. PAF itself may even represent a promising intervention target in older adults with ASD. Several studies show that increasing PAF through neurofeedback, or repetitive transcranial magnetic stimulation, leads to cognitive gains in healthy adults ^84,85^ and improves memory in participants with mild cognitive impairment ^86^. Particularly relevant to the present findings, interventions that increase PAF through neurofeedback improve cognitive processing speed and executive function in elderly individuals, ^87^ suggesting that modulating oscillatory activity may provide an opportunity to ameliorate sources of additional age-related risk associated with ASD.

The current study has several limitations. First, cross-sectional designs preclude the analysis of individual trajectories of PAF decline and introduce individual differences that may be conflated with aging effects. Likewise, the truncated age range (< 70 years) of the current sample may capture only the early signs of functional aging in NT adults, possibly contributing to the attenuated relationship between age and PAF observed in NT participants (P=0.09). Finally, due to differences in behavioral measures across the three datasets, assays of cognitive function were limited in the present analyses. Together, these issues highlight a need for future longitudinal studies that include older participants and employ a more comprehensive range of cognitive assessments, including assays of executive functioning, a domain that is particularly affected by typical brain aging ^88,89^ and sensitive to cognitive declines in ASD. ^20,90^

In summary, patterns of PAF decline are atypical in adults with ASD, suggesting that neurophysiological aging may be altered in ASD. In addition to highlighting a need for more research into underlying neurobiological aging mechanisms, the present results also indicate that neural markers may help identify individuals at risk of poor aging outcomes before behavioral changes emerge. Specifically, neurophysiological markers such as PAF may help us to identify risk factors throughout the life course that shape and define aging trajectories in later life. Ultimately, understanding aging processes and related risk factors at an individual level may inform preventive interventions to promote successful aging and improve quality of life for adults with ASD in our increasingly elderly society.

## Acknowledgement

Data and/or research tools used in the preparation of this manuscript were obtained from the National Institute of Mental Health (NIMH) Data Archive (NDA). NDA is a collaborative informatics system created by the National Institutes of Health to provide a national resource to support and accelerate research in mental health. Dataset identifier(s): 10.15154/1519182. This manuscript reflects the views of the authors and may not reflect the opinions or views of the NIH or of the Submitters submitting original data to NDA.

## Author Contributions

Abigail Dickinson contributed to the conception and design of the study, analysis of the data and drafting a significant portion of the manuscript. Shafali Jeste contributed to the conception of the study and drafted a significant portion of the manuscript. Elizabeth Milne contributed to the conception and design of the study, data acquisition, and drafted a significant portion of the manuscript.

