## Supplementary Materials for "Electrophysiological signatures of brain aging in autism spectrum disorder"

**Supplementary Information**

*Description of NDAR Participants*

Participants were included in the present study if they were (1) aged over 18 years, (2) had a confirmed ASD diagnosis, and (3) had successfully contributed resting-state EEG data. These criteria yielded N=65 participants with ASD collected as part of two independent studies. Age-matched NT control participants from these two studies (N=59) were also included in analyses. [ADD TABLES].

*ASD Assessments*

Six participants with ASD scored <60 on the SRS. These participants all scored above the threshold for ASD on the revised Ritvo Autism and Asperger Diagnostic Scale (RAADS-R) (Ritvo et al. 2011). The RAADS-R is an 80-item questionnaire that was developed to assist clinicians diagnosing ASD in adults. Possible scores range from 0-240, with higher scores indicating a higher level of ASD symptomatology. A score above 65 is consistent with an ASD diagnosis (Ritvo et al. 2011), whereas Anderson and colleagues (2011) recommend a more conservative cut-off score of 72 to maximize sensitivity and specificity (Andersen et al. 2011). All six participants who scored <60 on the SRS scored >74 on the RAADS-R. These participants were therefore retained in analyses, as they had received a clinical diagnosis and obtained a RAADS-R score above the most-conservative cut-off for ASD.

*EEG Data Harmonization*

Pooling EEG data from three independent studies required several study differences to be accounted for. We have described the main differences between EEG acquisition, and how we accounted for these differences, below.

Electrode Montage: EEG data were recorded from 64 channels in studies 1 &2, and 123 electrodes in study 3. All three montages included electrodes that represented standard locations in the 10-20 EEG system. We down-sampled the data to 25 of these standard electrode sites in order to use an equal number of comparable electrodes across all three studies.

Sampling Rate: Data were down-sampled to 512Hz for the present analyses. Using a common sampling rate across all three studies circumvented different sampling rates that were used during recording 20148Hz in study 1; 1000Hz in studies 2 & 3).

Impedances: There is no way to account for the variability in system impedances post-hoc. As electrode impedance describes the connection between the electrode and skin, higher impedances are associated with smaller EEG signal amplitude. However, several studies have now shown that the differences in impedances between research EEG systems do not affect the size of the EEG signal (Johnson et al. 2001; Kappenman and Luck 2010). It should also be noted that calculating peak alpha frequency is independent of EEG amplitude.

Reference: To account for reference differences between the three studies, all data were re-referenced to the average of all electrodes (after data had been cleaned and down-sampled to a consistent 25-channel montage).

Recording Length: Only the first 2 minutes of data were analyzed for each participant, meaning that differences in recording length across the three studies were not represented in the present analyses.

**Supplementary Tables**

Table S1. Participant characteristics for each study.

|  |  | | | | Study 1 | |  | Study 2 | |  | Study 3 | |
| --- | --- | --- | --- | --- | --- | --- | --- | --- | --- | --- | --- | --- |
|  |  | | | | ASD  N=28 | NT  N=28 |  | ASD  N=29 | NT  N=34 |  | ASD  N= 36 | NT  N=25 |
| Age (Years) | | | | |  |  |  |  |  |  |  |  |
|  | | Mean  (SD) | | | 44.34  (13.31) | 39.70  (10.97) |  | 26.19 (11.44) | 31.15 (14.37) |  | 22.93  (4.52) | 26.34  (4.54) |
|  | | Range | | | 18.08 – 67.42 | 24.67 – 68.33 |  | 18-64.58 | 19-69 |  | 18.17 – 35.75 | 18.42 – 26.34 |
| Sex | | | | |  |  |  |  |  |  |  |  |
|  | N females  (% Female) | | | | 12  (42.9%) | 15  (53.6%) |  | 5  (17.2%) | 15 (44.1%) |  | 9  (25%) | 4  (16%) |
| NVIQ | | | | |  |  |  |  |  |  |  |  |
|  | | *n* completed | | | 25 | 28 |  | 28 | 32 |  | 34 | 25 |
|  | | Mean  (SD) | | | 59.4 (  8.18) | 58.57  (7.09) |  | 52.29 (8.60) | 56.41 (11.77) |  | 48.17  (8.98) | 56.54  (7.59) |
|  | | Range | | | 29-67 | 49 - 75 |  | 40 - 70 | 33 – 77 |  | 30.50 - 68 | 36.5 – 69.5 |
| SRS | | | | |  |  |  |  |  |  |  |  |
|  | | | *n* completed | | 28 | 28 |  | - | - |  | - | 24 |
|  | | | Mean  (SD) | | 67.71  (10.01) | 46  (5.11) |  | - | - |  | - | 44.54  (6.18) |
|  | | | Range | | 48 - 84 | 39 - 57 |  | - | - |  | - | 38 - 62 |
| ADOS | | | | |  |  |  |  |  |  |  |  |
|  | | | | *n* completed | - | - |  | 29 | 34 |  | 35 | - |
|  | | | | Mean (SD) | - | - |  | 9.97  (3.41) | 2.18 (2.21) |  | 11.15  (3.51) | - |
|  | | | | Range | - | - |  | 4 - 19 | 0 - 7 |  | 7-21 | - |

Table S2. *Additional diagnosis and medication information for all participants.*

|  | **ASD** | **NT** |
| --- | --- | --- |
| **Additional Diagnoses** | | |
| Participants with a clinical diagnosis | 43% (40) | 5.7% (5) |
| Number of additional diagnoses | One: 33.3% (31)  Two: 9.7% (9) | One: 5.7% (5)  Two: 0% (0) |
| Conditions | Depression (11), Anxiety (7), ADHD (8), OCD (2), Bipolar Disorder (2), Depression & Anxiety (4), Depression & ADHD (2), ADHD & Anxiety (3), ADHD & OCD (1). | Depression (3), PTSD (1), Anxiety (1) |
| **Medications** | | |
| Participants taking medication | 30.1% (28) | 3.4% (3) |
| Number of medications | One: 22.6% (21)  Two: 7.5% (7) | One: 3.4% (3) |
| Medication Type | SSRI (10), AED (4), Stimulant (2), Tricyclic Antidepressant (1), Atypical Antipsychotic (1), NDRI (1), Benzodiazepine (1),  SSRI & AED (1), SSRI & Stimulant (2), SSRI & Sedative (1), SSRI & Atypical Antipsychotic (1), SSRI & Beta Blocker (1), SSRI & Benzodiazepine (1). | SSRI (1), antimetabolite (1). |

*In the ASD group, conditions additional to ASD diagnosis are listed.

Abbreviations: ADHD = attention deficit hyperactivity disorder, OCD= obsessive compulsive disorder, PTSD = post-traumatic stress disorder, SSRI= selective serotonin reuptake inhibitor, NDRI = Norepinephrine and dopamine reuptake inhibitor, AED = antiepileptic drug.

Table S3. *Hierarchical linear regression analyses of age-related change in average PAF.*

|  |  | *F* | *F* Change | *R^2^* | Intercept | Beta (*SD*) |
| --- | --- | --- | --- | --- | --- | --- |
| Baseline Model | | 1.59 | - | 0.03 | 9.28 |  |
|  | Sex |  |  |  |  | 0.08 (0.08) |
|  | Study |  |  |  |  | -0.03 (0.13) |
|  | NVIQ |  |  |  |  | **0.014 (0.01)*** |
| Second Block | | 6.09*** | 19.12*** | 0.13 | 10.41 |  |
|  | Age |  |  |  |  | **-0.02 (0.07)***** |
| Third Block | | 4.99*** | 0.64 | 0.13 | 10.41 |  |
|  | Group |  |  |  |  | -0.09 (0.12) |
| Fourth Block | | 5.43*** | 6.77* | 0.17 | 10.65 |  |
|  | Age by Group Interaction |  |  |  |  | **0.020 (0.01)*** |

Note. Significant predictors are denoted in bold. .**p*<0.05, ***p*<0.001, ****p*<0.0001
